# Therapeutic strategy for spinal muscular atrophy by combining gene supplementation and genome editing

**DOI:** 10.1101/2023.04.06.535786

**Authors:** Fumiyuki Hatanaka, Keiichiro Suzuki, Kensaku Shojima, Jingting Yu, Yuta Takahashi, Akihisa Sakamoto, Javier Prieto, Maxim Shokhirev, Concepcion Rodriguez Esteban, Estrella Nuñez-Delicado, Juan Carlos Izpisua Belmonte

## Abstract

Defect in the *SMN1* gene causes spinal muscular atrophy (SMA), which shows loss of motor nerve cells, muscle weakness and atrophy. While current treatment strategies, including small molecules or viral vectors, have been reported to improve motor function and survival, an ultimate and long-term treatment to correct SMA endogenous mutations and improve its phenotypes is still highly challenging. We have previously developed a CRISPR-Cas9 based homology-independent targeted integration (HITI) strategy, which allowed for unidirectional DNA knock-in in both dividing and non-dividing cells *in vivo*. Here, we demonstrated its utility by correcting a SMA mutation in mice, and when combined with *Smn1* cDNA supplementation show SMA long-term therapeutic benefits in mice. Our observations may provide new avenues for long term and efficient treatment of inherited diseases.

**Summary:** The Gene-DUET strategy by combining cDNA supplementation and genome editing was sufficient to ameliorate SMA phenotypes in mouse model *in vivo*.

## INTRODUCTION

Spinal muscular atrophy (SMA) is a severe inherited neuromuscular disease caused by deletion or mutation of survival motor neuron gene 1 (*SMN1*), which contributes to numerous cellular processes and pathways, the most studied function is its role in snRNP assembly*(1)*. SMA is characterized by the degeneration of lower motor neurons which leads to muscle weakness and atrophy. SMA occurs in approximately 1 in 11,000 newborns and represents the most common genetic cause of infant mortality*(2)*.In human, there are two forms of SMNs. *SMN1* is the primary gene for production of SMN protein. *SMN2* is a paralog of *SMN1*, which differs only a few nucleotides and generates a truncated unstable protein with low levels of full-length SMN protein compared as *SMN1*. Although only 10-20% of the *SMN2* gene product is fully functional, it can partially compensate for the loss of *SMN1* and increased copy number of *SMN2* is inversely correlated with disease severity in SMA patients*(3)*. Today, there are several approaches to treat SMA and effectively prolong the life of patients. Antisense oligonucleotides (ASOs) targeted to RNA splicing of *SMN2* gene produces a full-length mRNA and a functional SMN protein*(4)*. ASOs have huge potential in SMA therapy but present some limitations including the difficulties crossing the blood-brain barrier. Another alternative approach for SMA is gene supplementation therapy, which delivers a fully functional copy of the *SMN1* gene on the genome of self-complementary adeno-associated viral (AAV) vector packaged with serotype 9 (scAAV9). Recently, Mendell et al. showed that an one-time intravenous injection of high dose scAAV9-*SMN1* resulted in improved motor function and extended survival in SMA patients*(5)*. However, this gene supplementation therapy cannot permanently restore endogenous *SMN1* expression because the genome of the recombinant AAV vectors does not undergo genomic integration into the host genomic DNA. The AAV genomes persist mainly in an episomal form in the nucleus of the host cells, which eventually leads to dilution of the gene encoded by the AAV genome. Therefore, new strategies for *in situ* correction of endogenous mutated sequences are needed for efficient and long-term improvement of SMA.

The CRISPR-Cas9 technology is a powerful genome-editing system which could be applied to potential cure for common inherited diseases*(6, 7*). However, *in situ* gene correction of the SMA-causing mutation has not been reported, likely due to the inherent difficulties in accessing motor neurons within the spinal cord. A major hurdle is that genome editing in non-dividing cells, including motor neurons, is inefficient in the absence of the homology-directed repair (HDR) pathway used in conventional gene-repair approaches. Another problem is the low *in vivo* delivery of genome-editing tools into the spinal cord, which prevents sufficient genome correction to improve the disease phenotypes.

We recently developed a non-homologous end joining (NHEJ)-based homology-independent strategy for targeted integration (HITI) of transgenes in both dividing and non-dividing cells*(6, 9*). Since NHEJ is active throughout the cell cycle in a variety of cell types including neurons*(6, 11*), the HITI technology has the potential to provide a solution to some of the current challenges for efficient and long-term amelioration of SMA phenotypes. In here, by combining gene cDNA supplementation and genome editing using HITI (from now on, Gene-DUET), we show long-term survival and amelioration of SMA phenotypes. Beside improving current therapies for SMA, our observations may have implications for the treatment of other inherited diseases especially for neurodegenerative disorders as well as neuromuscular diseases.

## RESULTS

### AAV-mediated *in vivo* genome editing in spinal cord

AAV vector has emerged as one of the safest and most commonly used vectors for the delivery of therapeutic genes*(12)*. In fact, AAVs have been used extensively in gene therapy clinical trials to treat patients with SMA, Duchenne muscular dystrophy (DMD) and X-linked myotubular myopathy patients *(6, 12, 13*).Over the past few decades, numerous engineered AAV capsids have been developed for effective gene delivery into various tissues, and current strategies for developing AAV vectors with tailored tropism are described*(14)*. Recent report showed that engineered AAV-PHP.eB capsid enables efficient transduction in the central nervous system*(15)*. Then, we first compared AAV-PHP.eB and AAV9 which is commonly used for clinical gene therapy of SMA patients. We injected GFP-expressing AAV (AAV-GFP) into neonatal wild-type (WT) mouse by intravenous injection and analyzed GFP expression in the brain, lung, heart, stomach, liver, spleen, pancreas, kidney, muscle and spinal cord 2 weeks later (Fig. 1, A and B, and fig. S1, A and B). The analyses of tissue sections showed that AAV-PHP.eB was more efficient in the spinal cord and brain than AAV9 (fig. S1, C to F). Notably, the GFP signals were abundantly merged with NeuN which is a motor neural marker in the spinal cord (Fig. 1C). These results demonstrate the advantage of AAV-PHP.eB for transduction into spinal motor neurons.

**Figure 1.**
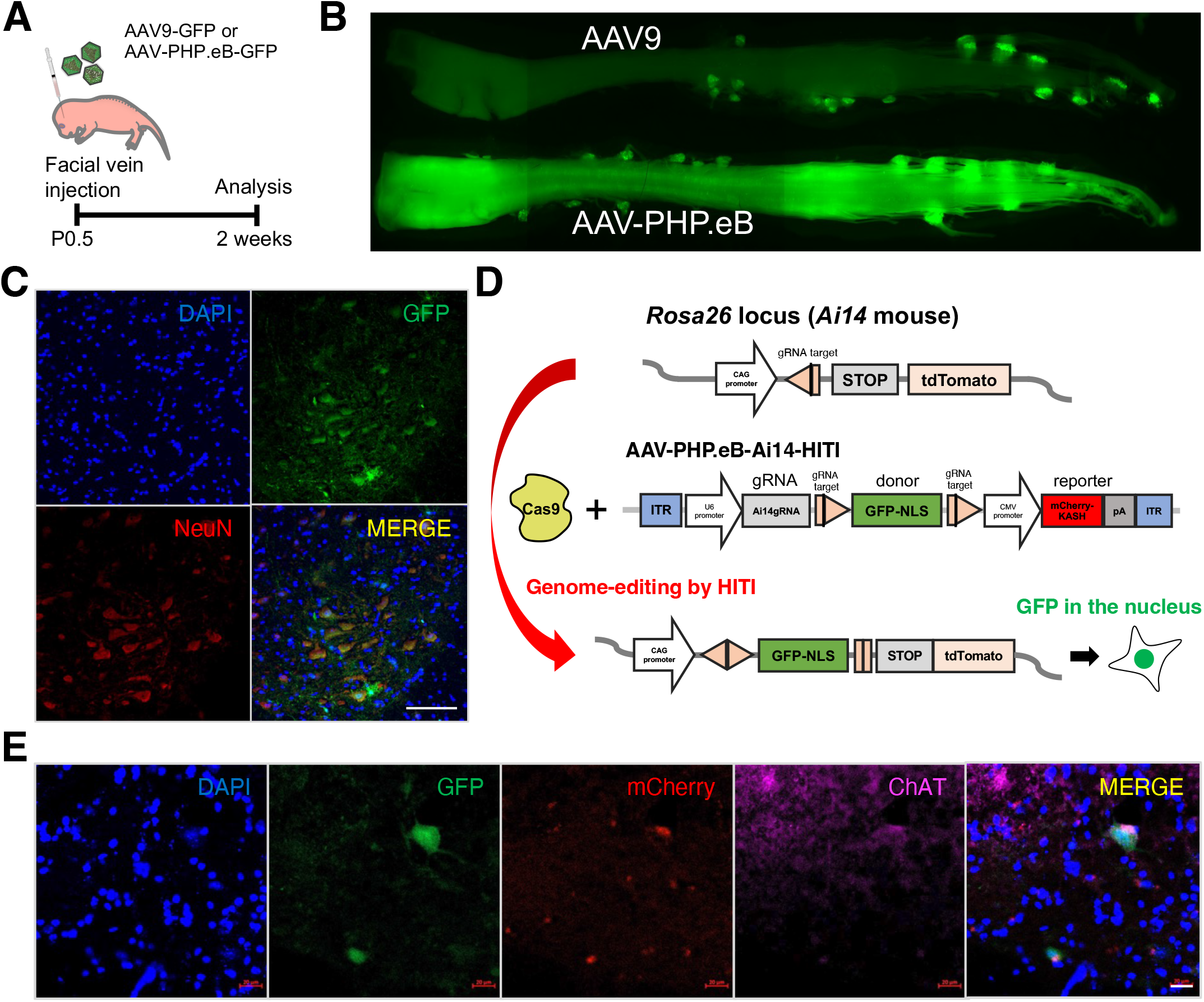

To evaluate the frequency of targeted gene knock-in *in vivo*, we previously used Ai14 mice carrying the CAG promoter at the *Rosa26* loci (Fig. 1D)*(8)*. Since the HITI technology can perform *in vivo* gene knock-in in non-dividing cells represented by neural cells, we tested the *in vivo* efficacy of HITI in the spinal cord by using Ai14-Cas9 mice which constitutively express Cas9. HITI-mediated GFP knock-in at the *Rosa26* locus downstream of the CAG promoter was observed in the nucleus of the motor neurons and liver when the AAV-PHP.eB-Ai14-HITI including guide RNA (gRNA) expression cassette, GFPNLS-pA donor sandwiched by Cas9/gRNA target sequence and mCherry reporter were delivered into Ai14-Cas9 mice at postnatal day 0.5 (P0.5) (Fig. 1E and fig. S1, G and H). These results suggest that HITI-mediated genome editing is successful in the spinal motor neurons.

### Targeted *in vivo* gene correction of SMA mice via HITI

Next, to demonstrate the validity of the HITI technology for gene correction of SMA, we chose the SMA mouse (*SMN2*^+/+^; SMNΔ7^+/+^; *Smn1*^−/−^) as a disease model, which disrupts endogenous *Smn1* gene (*mSmn1*) through lacZ reporter gene insertion into exon 2 and harbor two transgenic alleles of human *SMN2(16–18)*. To prevent the deleterious effect of CRISPR-Cas9-induced insertions/deletions (indels) in the endogenous exon, we targeted intronic sequences upstream of the exon2/lacZ in chromosome 13 (Fig. 2A). Previously, we demonstrated that unknown recombination occurs within homologous sequence between genome and donor*(19)*. To avoid such an unexpected event, we removed the homologous sequence on the donor by incorporating a portion of rat intron 1 of *Smn1* including the splicing acceptor, codon-optimized mouse *Smn1* cDNA (exons 2-8), and rat 3’UTR. The constructed pAAV-*SMN1*-HITI vector contains intron 1 targeting gRNA expression cassette and *SMN1* donor sandwiched by two gRNA target sequences. We packaged the pAAV-*SMN1*-HITI and an AAV expressing Cas9 (pAAV-Cas9) with PHP.eB capsid, and systemically delivered the AAV-PHP.eB-*SMN1*-HITI and AAV-PHP.eB-Cas9 via intravenous injection into SMA mice at P0.5 (Fig. 2B). HITI-mediated gene knock-in was detected by PCR amplification only in treated tissue 2 weeks after injection (Fig. 2C). We also verified the corrected genome sequences in amplicons by Sanger sequencing (fig. S2A). Importantly, HITI-treated SMA mice showed phenotypic improvement compared to untreated SMA mice. HITI-treated SMA mice were able to walk independently whereas untreated SMA mice were unable to stand at 2 weeks old (Fig. 2D and Movie S, 1 and 2). The mean survival and body weight were significantly increased in HITI-treated SMA mice than in untreated SMA mice (Fig. 2, E and F, and fig. S2B). However, the effect of HITI was not enough for SMA mice to survive more than 3 weeks even showing significant differences in behavior and survival analyses. These results suggest that HITI-mediated genome editing is successful in SMA mice and rescues SMA phenotypes, but is not sufficient for a therapeutic strategy for SMA.

**Figure 2.**
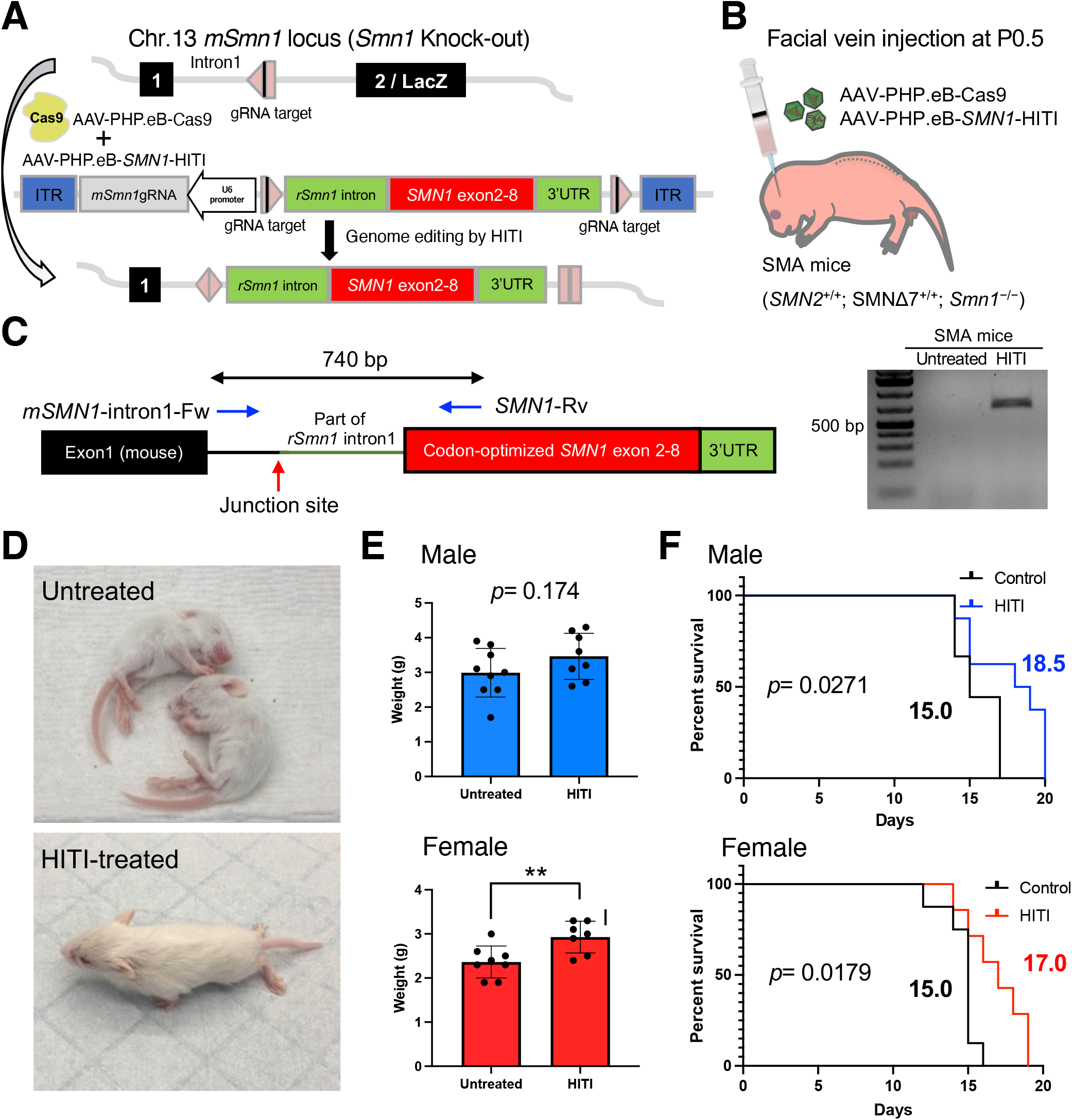

### Gene-DUET strategy for SMA through gene supplementation and genome editing

The body weight of SMA mice at birth was lower than that of WT and heterozygous litters, suggesting that SMA phenotypes were already advanced at the time of treatment (fig. S2C). HITI-mediated gene correction by AAV and the resulting SMN1 protein production is likely to take a several days, which is too late to rescue SMA mice effectively. Previous reports also showed that the timing of initial treatment and high SMN1 expression was important for SMA therapy*(6, 21*). Based on these results, we established a new strategy, Gene-DUET, which is a combination of two strategies through wildtype cDNA supplementation and genome editing. Wild-type cDNA supplementation was accomplished by overexpressing mouse *Smn1* (*mSmn1*) cDNA as previous reports*(22)*. We redesigned the AAV vector (AAV-*SMN1*-DUET) containing the *mSmn1* coding sequence (CDS) under the CMV promoter and *SMN1*-HITI as used in the previous construct in Fig. 2A. Similar to current gene supplementation therapy, only *mSmn1* CDS would be expressed in the absence of Cas9 (Fig. 3A). In contrast, *mSmn1* CDS and the gene-corrected *mSmn1*/*SMN1* fusion gene can be co-expressed in the presence of Cas9 (Fig. 3B). We systemically delivered AAV-PHP.eB-*SMN1*-DUET with or without AAV-PHP.eB-Cas9 via intravenous injection into SMA mice at P0.5 (Fig. 3, C and D). The cDNA alone (without Cas9) or DUET (with Cas9) treated SMA mice looked very healthy compared to untreated SMA mice in general appearance 2 weeks after the injections (Fig. 2D, 3E and Movie S, 3 and 4). Dissection of tissues at 2 weeks showed that cDNA or DUET treatments improved the size of the spinal cord, brain, heart and muscle (Fig. 3, F and G, and fig. S3, A to E). We evaluated motor function in each treated SMA mice by analyzing the righting reflex test 2 weeks after the injections. Untreated WT and heterozygous *Smn1* mice were able to right themselves quickly, whereas the SMA mice took longer or were unable to right themselves within 30 seconds. The cDNA-or DUET-treated SMA mice significantly showed improvement of righting reflex compared to untreated SMA mice in both male and female mice (Fig. 3H). Some of the treated SMA mice exhibited ear and digital necrosis and shorter tails due to necrosis starting at 5 weeks of age until loss as reported in previous reports (fig. S3F)*(23, 24*). These results suggest that both cDNA and DUET treatments dramatically improve the phenotypes of SMA mice.

**Figure 3.**
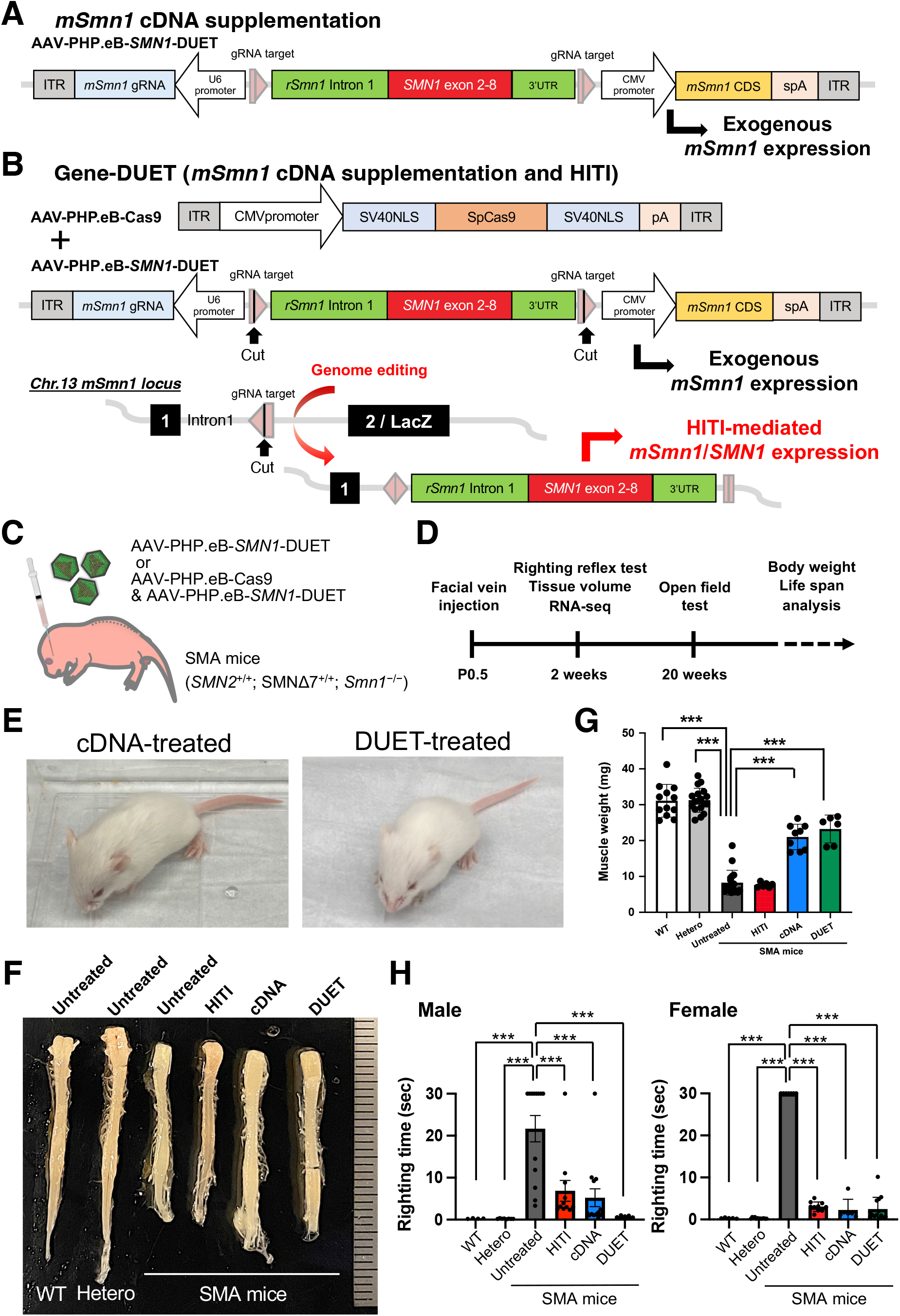

### Molecular restoration in SMA mice by cDNA and DUET treatments

We next performed RNA sequencing (RNA-seq) of spinal cord samples to understand the global transcriptional alterations by HITI, cDNA and DUET treatments in SMA mice compared to untreated SMA mice and control heterozygous mice. Principal component analysis (PCA) and heatmap analyses clearly segregated untreated SMA mice from the healthy heterozygous mice (Fig. 4A and fig. S4A). The profile of cDNA- and DUET-treated SMA samples was placed close to that of heterozygous mice, suggesting restoration by treatments against SMA-induced molecular dysfunction. A functional gene enrichment analysis revealed that inflammatory pathways including p53 signaling pathway and cytokine-cytokine receptor interaction were up-regulated while motor neuron pathway including cholinergic synapse was down-regulated in SMA mice compared to heterozygous mice (Fig. 4, B and C, and fig. S4, B and C). These molecular changes were restored by both cDNA- and DUET-treatments but not restored by HITI treatment. (Fig. 4D). We performed quantitative reverse transcription PCR (RT-qPCR) and confirmed that activated p53 downstream genes including *Gtse1, Ccng1, Perp* and *Sesn1* were significantly repressed in cDNA- and DUET-treated spinal cord compared to untreated SMA samples (fig. S4D). These results suggest that both cDNA- and DUET-treatments restore the molecular dysfunction in the spinal cord of SMA mice.

**Figure 4.**
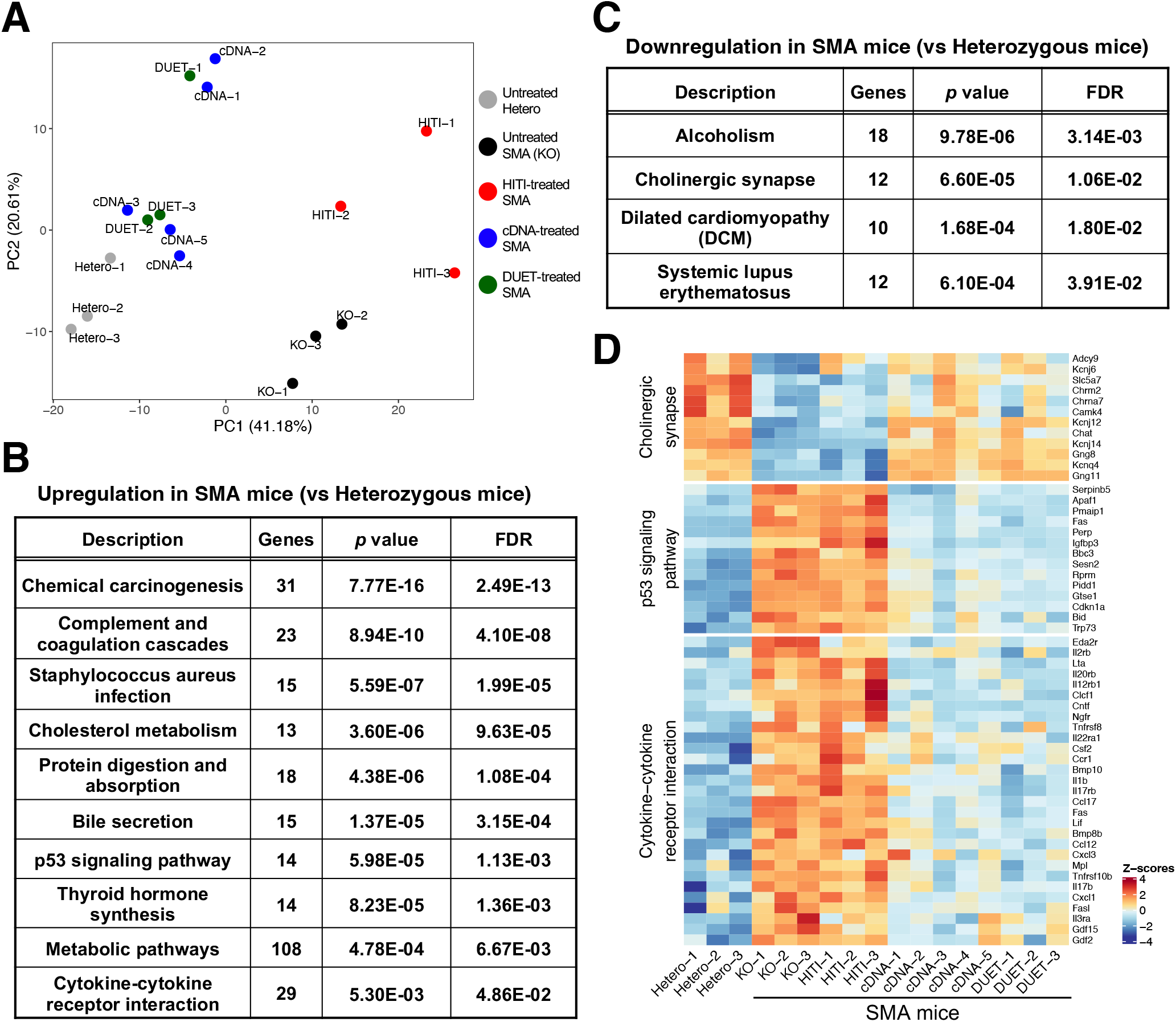

### Stable gene correction mediated by Gene-DUET

General appearance at 20 weeks old was not different between cDNA- and DUET-treated SMA mice (Fig. 5A). In terms of body weight, cDNA-or DUET-treated SMA mice showed dramatic improvement compared to untreated SMA mice (Fig. 5B). Locomotor activity of cDNA- and DUET-treated SMA mice at 20 weeks old was similar to that of heterozygous mice, indicating that motor function was significantly improved even into adulthood (fig. S5). The concern with neonatal AAV treatment is the loss of the transgene along with cell division during tissue growth. Previous report showed quick reduction of vector genome copies in the liver over a few weeks after neonatal AAV treatment*(25)*. Indeed, GFP expression in the spinal cord was declined after 1 year compared to 2 weeks in AAV-PHP.eB-GFP injected WT mice (Fig. 1B, and fig. S6A). We also found a significant reduction in cDNA-derived exogenous *mSmn1* expression in the spinal cord and liver at 1 year old compared to at 2 weeks old in cDNA-treated heterozygous mice (fig. S6, B and C). Theoretically, we thought that genome editing by HITI might be more advantageous than cDNA supplementation for permanent cure. To examine the efficiency of gene correction in DUET-treated SMA mice, we extracted genomic DNA from the treated mice and enriched the targeting regions by using customized probes that could hybridize and pull down the genomes around the Cas9 cleavage site (Fig. 5C). We performed the deep sequencing and calculated the editing efficiency in 20- and 40-weeks old DUET-treated tissues. The corrected sequences were detectable and stable in all tissues from these DUET-treated SMA mice (Fig. 5, D and E). These results suggest that gene correction by Gene-DUET strategy is stable for a long time in SMA mice. Crucially, both cDNA and DUET treatments significantly enhanced the survival of SMA mice compared to untreated SMA mice. More importantly, the mean survival of DUET-treated SMA mice was clearly improved over that of cDNA-treated SMA mice, especially in males (Fig. 5, F and G). These data suggest the synergistic effect of the Gene-DUET strategy by gene supplementation and genome editing.

**Figure 5.**
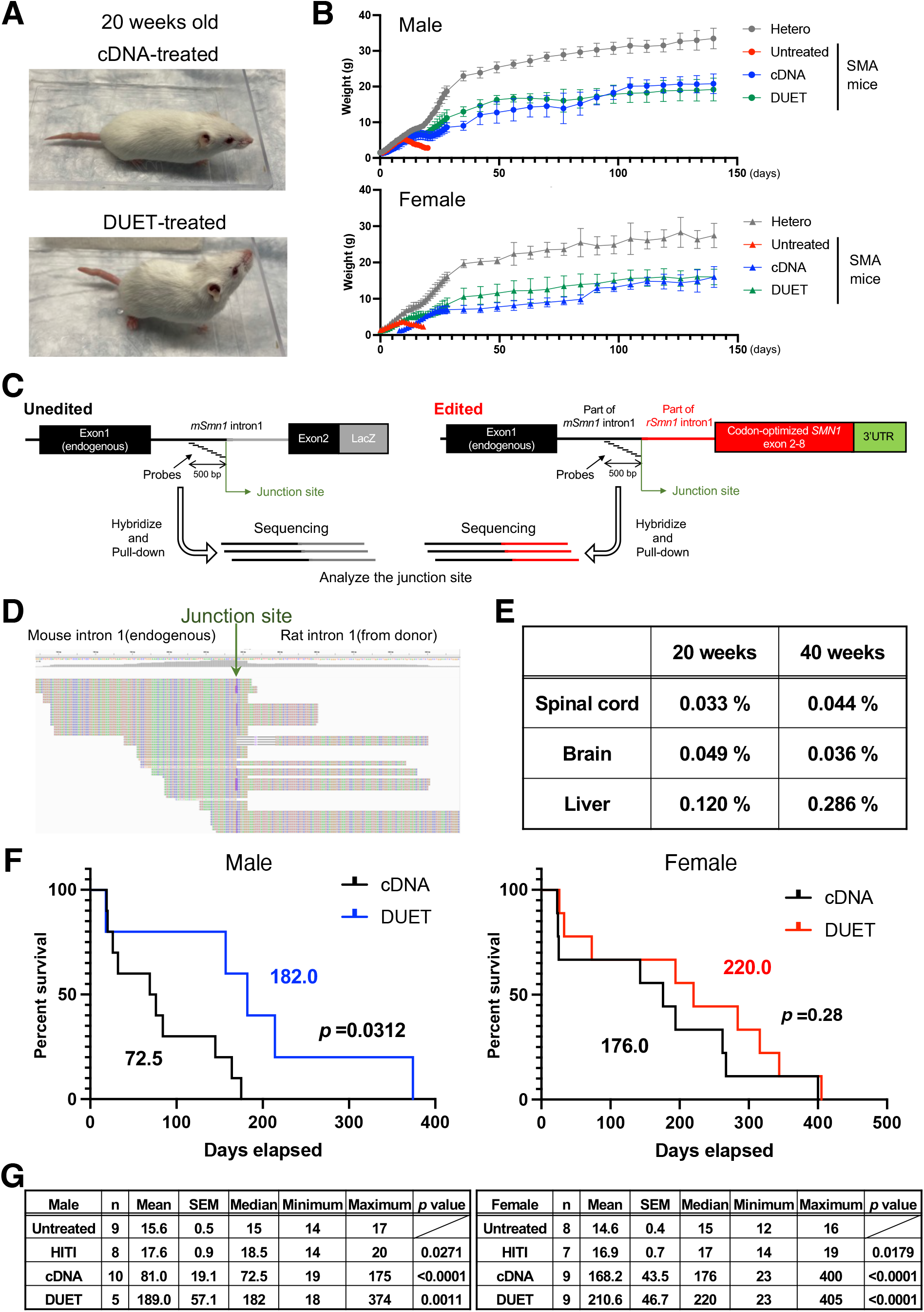

## DISCUSSION

Gene therapy medicine, Zolgensma, for treating SMA by cDNA supplementation has been recently approved by FDA. However, the long-term efficacy of this treatment has not yet been determined. Indeed, our data show that exogenous *Smn1* expression is attenuated in spinal cord and liver cells one year after AAV injection. The development of HITI technology has enabled the correction of genomic mutations in non-dividing cells through the NHEJ pathway and here we provide a first demonstration of its utility in an SMA model in mice. Compared to cDNA supplementation, the *in situ* gene correction approach here reported is able to sustain stable and permanent *Smn1* expression. However, due to the widespread and profound phenotypic alterations that arise immediately after birth, gene correction alone was not sufficient to correct all of them although it is able to correct a subset of some of the phenotypic changes observed in SMA mice. The extent of cell and tissue phenotypic restoration and increase in survival time were observed when combining HITI-mediated genome editing and gene supplementation in SMA mice. We observed a more significant survival benefit in male mice than in female mice by Gene-DUET. This is likely due to sex differences in SMA phenotypes and tissue growth differences between male and female mice, with exogenous *mSmn1* being retained more in female than in male mice*(26)*. Previous report showed that sex-specific amelioration of SMA phenotype by an antisense oligonucleotide treatment*(27)*. In fact, we also observed that the mean survival of cDNA-treated female SMA mice was two times longer than that of cDNA-treated male SMA mice.

Recently, the base editing combined with antisense oligonucleotide can ameliorate the SMA phenotypes*(28)*, however that therapeutic effects depend on the copy number of endogenous *SMN2*, which may not be sufficient for SMA type 0 and type 1 patients. As we have shown in this work, the Gene-DUET supports therapeutic benefit for SMA mice compared to conventional gene supplementation therapy. In summary, our Gene-DUET strategy provides new exploratory avenues for the treatment of SMA in humans. This approach has a great potential for the field of genome-editing technologies that may be relevant for the treatment of inherited diseases, particularly neurodegenerative and neuromuscular diseases.

## MATERIAL AND METHODS

### Plasmids

pAAV-CAG-GFP (addgene 37825) and pUCmini-iCAP-PHP.eB (addgene 103005) were purchased from Addgene. pAAV-CMVc-Cas9 (addgene 106431) was established previously*(29)*. To construct gRNA expression vectors, each 20 bp target sequence was sub-cloned into pCAGmCherry-gRNA (Addgene 87110). The CRISPR/Cas9 target sequences (20 bp target and 3 bp PAM sequence (underlined)) used in this study are shown as following: *Smn1* intron 1 targeting gRNA (GCCATACCATAAGACGACCGAGG). The downstream of CAG promoter in Ai14 mouse (TAGGAACTTCTTAGGGCCCGCGG) gRNA expression plasmid has been established in the previous paper *(8)*. pAAV-nEFCas9 (Addgene 87115) and AAV-Ai14-HITI (Addgene 87117) were established in the previous paper*(8)*. To construct donor/gRNA AAV for HITI-mediated *Smn1* gene correction, part of intron 1 including splicing acceptor site, 3’UTR and downstream were amplified from rat genome isolated from Brown Norway rat. The mouse *Smn1* exon 2-8 was codon optimized and synthesized in IDT. The assembled fragment was sandwiched by two *Smn1* intron 1 gRNA target sequence and subcloned into between ITRs of PX552 purchased from Addgene (Addgene 60958), and generated pAAV-*SMN1*-HITI. To construct pAAV-*SMN1*-DUET, CMV-*mSmn1* CDS expression cassette (ORIGENE MR203917) was sub-cloned into pAAV-*SMN1*-HITI.

### Animals

C57BL/6, Rosa-LSL-tdTomato (known as Ai14), Rosa26-Cas9 knock-in mice (Stock No: 024858), SMNΔ7 mice (FVB.Cg-*Grm7*^*Tg(SMN2)89Ahmb*^ *Smn1*^*tm1Msd*^ Tg(*SMN2*\*delta7)4299Ahmb/J, Stock No: 005025) were purchased from the Jackson laboratory. We mated Ai14 mouse and Rosa26-Cas9 knock-in mouse to generate the Ai14-Cas9 mouse. The mice were housed in a 12-hours light/dark cycle (light between 06:00 and 18:00) in a temperature-controlled room (22 ± 1 °C) with free access to water and food. All procedures were performed in accordance with protocols approved by the IACUC and Animal Resources Department of the Salk Institute for Biological Studies.

### AAV production

AAV9 and AAV-PHP.eB viral particles were generated by or following the procedures of the Gene Transfer Targeting and Therapeutics Core at the Salk Institute for Biological Studies.

### Facial vein AAV injection

The newborn (P0.5) mice were used for intravenous AAV injection as following previous report*(29)*. The AAV mixtures were injected via temporal vein of the newborn mice. After the injection, pups were allowed to completely recover on a warming pad and then returned to the home cage.

### Intraspinal AAV injection

Neonatal pups were used for spinal cord AAV injection as following previous report*(30)*. Briefly, P0.5 mice were anesthetized and 2 μl of AAV mixtures was slowly injected into the spinal cord using 33 Gauge Neuros syringe (65460-06, Hamilton,). After the injection, pups were allowed to completely recover on a warming pad and then returned to the home cage.

### Genome extraction

Genomic DNA was extracted from cells and tissue samples using DNeasy Blood & Tissue Kits (69506, Qiagen) following the manufacturer’s instruction.

### RNA Analysis

Total RNA was extracted from cells and tissue samples using either TRIzol (Invitrogen) or RNeasy Kit (Qiagen) followed by cDNA synthesis using Maxima H Minus cDNA Synthesis Master Mix (Thermo Fisher). Real-time qPCR was performed using SsoAdvanced SYBR Green Supermix and analyzed using a CFX384 Real-Time system (Bio-Rad). All analyses were normalized based on amplification of mouse *Gapdh*.

### Immunohistochemistry

Tissues were harvested after transcardial perfusion using ice-cold PBS, followed by ice-cold 4% paraformaldehyde in phosphate buffer for 15 min. Tissues were dissected out and postfixed in 4% paraformaldehyde overnight at 4 °C and cryoprotected in 30% sucrose overnight at 4 °C and embedded in OCT (Sakura Tissue-Tek) and frozen on dry ice. Serial sections at 12 μm were made with a cryostat and collected on Superfrost Plus slides (Fisher Scientific) and stored at −80 °C until use. Immunohistochemistry was performed as follows: sections were washed with PBS for 5 min 3 times, incubated with a blocking solution (PBS containing 2% donkey serum (or 5% BSA) and 0.3% Triton X-100) for 1 h, incubated with primary antibodies diluted in the blocking solution overnight at 4 °C, washed with PBST (0.2% Tween 20 in PBS) for 10 min 3 times, incubated with secondary antibodies conjugated to Alexa Fluor 488, Alexa Fluor 546, or Alexa Fluor 647 (Thermo Fisher) for 1 h at room temperature. After washing, the sections were mounted with mounting medium (DAPI Fluoromount-G, SouthernBiotech). The primary antibodies used in this study were anti-GFP, 1:500 (Aveslabs); anti-NeuN, 1:100 (MCA-1B7, EnCor Biotechnology); anti-ChAT, 1:200 (NB110-89724, Novus Biologicals).

### Image Capture and Processing of Primary Neurons

Immunocytochemistry samples of mice samples were visualized by confocal microscopy using a Zeiss LSM 710 Laser Scanning Confocal Microscope (Zeiss). Images were processed by ZEN2 Black edition software (Zeiss).

### Righting reflex test

The righting reflex of untreated or treated mice was compared at postnatal 14 days. Mice were laid on their back and the time needed to flip over was recorded, with a maximum of 30 sec allowed. Three trials were performed for each mouse and the shortest time was used for analysis.

### Open field test

Mice were individually placed into clear Plexiglass boxes (40.6 × 40.6 × 38.1 cm) surrounded by multiple bands of photo beams and optical sensors that measure horizontal (ambulatory) and vertical (rearing) activity (Med Associates, USA). Mice movement was detected as breaks within the beam matrices and automatically recorded for 30 min.

### RNA sequencing and Data analysis

RNA was extracted from the spinal cord and prepared for RNA sequencing with TruSeq Standard mRNA sample Prep Kit (Illumina). Deep sequencing was performed on the Illumina NovaSeq6000, at pair end 150 bp. Sequenced reads were quality-tested using FASTQC v0.11.8 (https://www.bioinfomatics.babraham.ac.uk/projects/fastqc/) and adapters were trimmed using Cutadapt v2.4*(31)* with parameters ‘-j 8 -f fastq -e 0.1 -q 20 -O 1 -a AGATCGGAAGAGC’. Paired reads after trimming were aligned to custom mouse transcriptome and quantified at transcript level using Kallisto v0.46.0*(32)* with arguments ‘quant -b 30’. The custom transcriptome index was built with ‘kallisto index’ by merging the cDNA FASTA files of mm10 mouse transcriptomes and three genome-editing conditions of gene *Smn1* (including cDNA, HITI and Gene-DUET). Raw gene expression was estimated and summarized from transcript-level abundance with R package, tximport v1.12.3. Differential gene expression was performed on the raw gene counts with the R package, DESeq2 v1.24.0, using replicates to compute within-group dispersion. Differentially expressed genes were defined as having a false discovery rate (FDR) <0.05 and a |Fold Change| >1.5 when comparing two experimental conditions. Principle Component Analysis (PCA) was carried out on normalized gene counts using the R prcomp function. Heatmaps were generated with the R package, ComplexHeatmap v2.0.0. Overrepresentation Analysis (ORA) was performed with WebGestalt*(33)* using KEGG pathways with FDR <0.05 as the significance threshold, protein coding genes as the reference list, a minimum number of genes in a category of 5, and visualizing as a barplot with normalized enrichment scores.

### Target enrichment-genome sequencing analysis

Target enrichment-genome sequencing was performed using the SureSelect XT HS2 DNA system for Illumina Multiplexed Sequencing (Agilent Technologies) with a customized capture library following manufacture’s protocol. The customized probes to capture specific genomic loci were designed to include chr13:100125170-100125670 which 500bp is upstream of Cas9 cleavage site. 50 ng of genomic DNA was fragmented by using SureSelect Enzymatic Fragmentation Kit. The DNA libraries were prepared by using SureSelect XT HS2 DNA Library preparation Kit and amplified by unique dual indexing primer pair following manufacture’s protocol. After hybridization by customized probes, captured libraries were purified using streptavidin beads and amplified by PCR. Following indexing and sample pooling, sequencing was conducted using Illumina MiniSeq, at pair-end 150bp. Sequenced reads were trimmed using cutadapt v2.4*(31)* with parameters ‘-j 8 -f fastq -e 0.1 -q 20 -O 1 -a AGATCGGAAGAGC’ and then mapped to a custom genome reference with BWA v0.7.12*(34)* using default parameters. The custom reference consisted with DNA sequences of mouse chromosome 13 and the edit region around Cas9 cleavage site, upstream of which included the exon 1 and intron 1 of mouse *Smn1* and downstream of which included part of rat *SMN1* intron and optimized mouse exon 2-8. Resulting sam files were converted to sorted indexed bam files using samtools v1.9. Reads mapping to specific genomic region was obtained using ‘samtools view’ and then converted into bed format using bedtools v2.3 with arguments ‘bamToBed -cigar’, from which number of fully matched reads, reads with mismatch or indels were counted. Resulting counts were manually processed to produce position-specific figures and analyses.

## Statistical analysis

All of the data are presented as the mean ± S.D. or S.E.M and represent a minimum of three independent experiments. Statistical parameters including statistical analysis, statistical significance in the Figure legends. For statistical comparison, Two-tailed Student’s *t*-test and Log-rank (Mantel-Cox) test for survival curves were performed by Prism 9 Software (GraphPad).

## Data and software availability

The accession number for the RNA-seq data reported in this paper is NCBI GEO: GSE207181.

## List of Supplementary Materials

Fig. S1 to 6

## Acknowledgments

We are grateful to Grace Chou, Tzu-Wen Wang, Ling Ouyang and Nasun Hah for next-generation sequencing; Animal Resources Department for animal care; M. Schwarz and P. Schwarz for administrative help.

## Funding

The Uehara Memorial Foundation (FH, KShojima)

The Leave a Nest Research Grant (FH)

The Cell Science Research Foundation (KShojima)

Hashimoto Municipal Hospital Foundation (KShojima)

UCAM, Fundacion Dr. Pedro Guillen (JCIB)

The Japanese Society for the Promotion of Science KAKENHI, 21H04811 (KSuzuki)

AMED, JP22ek0109521 (KSuzuki)

## Author contributions

Conceptualization: FH, KSuzuki, JCIB

Methodology: FH, KSuzuki, KShojima, AS, JP, YT

Investigation: FH,

Bioinfomatics: JY, MNS

Project administration: CRE, END Writing – original draft: FH

Writing – review & editing: FH, KSuzuki, KShojima, JCIB

## Data and materials availability

Data supporting the findings of this study are available within the paper and its supplementary information files. RNA-sequence data was deposited in the Gene Expression Omnibus under the accession number GSE207181.

## References

1. H. Chaytow, Y.-T. Huang, T. H. Gillingwater, K. M. E. Faller, The role of survival motor neuron protein (SMN) in protein homeostasis. Cell Mol Life Sci 75, 3877–3894 (2018).

2. E. A. Sugarman, N. Nagan, H. Zhu, V. R. Akmaev, Z. Zhou, E. M. Rohlfs, K. Flynn, B. C. Hendrickson, T. Scholl, D. A. Sirko-Osadsa, B. A. Allitto, Pan-ethnic carrier screening and prenatal diagnosis for spinal muscular atrophy: clinical laboratory analysis of >72 400 specimens. Eur J Hum Genet 20, 27–32 (2012).

3. M. E. R. Butchbach, Copy Number Variations in the Survival Motor Neuron Genes: Implications for Spinal Muscular Atrophy and Other Neurodegenerative Diseases. Front Mol Biosci 3, 7 (2016).

4. Y. Hua, K. Sahashi, G. Hung, F. Rigo, M. A. Passini, C. F. Bennett, A. R. Krainer, Antisense correction of SMN2 splicing in the CNS rescues necrosis in a type III SMA mouse model. Genes Dev. 24, 1634–1644 (2010).

5. J. R. Mendell, S. Al-Zaidy, R. Shell, W. D. Arnold, L. R. Rodino-Klapac, T. W. Prior, L. Lowes, L. Alfano, K. Berry, K. Church, J. T. Kissel, S. Nagendran, J. L’Italien, D. M. Sproule, C. Wells, J. Cardenas, M. D. Heitzer, A. Kaspar, S. Corcoran, L. Braun, S. Likhite, C. Miranda, K. Meyer, K. D. Foust, A. H. M. Burghes, B. K. Kaspar, Single-Dose Gene-Replacement Therapy for Spinal Muscular Atrophy. New England Journal of Medicine 377, 1713–1722 (2017).

6. L. Cong, F. A. Ran, D. Cox, S. Lin, R. Barretto, N. Habib, P. D. Hsu, X. Wu, W. Jiang, L. A. Marraffini, F. Zhang, Multiplex Genome Engineering Using CRISPR/Cas Systems. Science 339, 819–823 (2013).

7. M. Jinek, K. Chylinski, I. Fonfara, M. Hauer, J. A. Doudna, E. Charpentier, A Programmable Dual-RNA–Guided DNA Endonuclease in Adaptive Bacterial Immunity. Science 337, 816–821 (2012).

8. K. Suzuki, Y. Tsunekawa, R. Hernandez-Benitez, J. Wu, J. Zhu, E. J. Kim, F. Hatanaka, M. Yamamoto, T. Araoka, Z. Li, M. Kurita, T. Hishida, M. Li, E. Aizawa, S. Guo, S. Chen, A. Goebl, R. D. Soligalla, J. Qu, T. Jiang, X. Fu, M. Jafari, C. R. Esteban, W. T. Berggren, J. Lajara, E. Nuñez-Delicado, P. Guillen, J. M. Campistol, F. Matsuzaki, G.-H. Liu, P. Magistretti, K. Zhang, E. M. Callaway, K. Zhang, J. C. I. Belmonte, In vivo genome editing via CRISPR/Cas9 mediated homology-independent targeted integration. Nature 540, 144–149 (2016).

9. K. Suzuki, J. C. I. Belmonte, In vivo genome editing via the HITI method as a tool for gene therapy. J Hum Genet 63, 157–164 (2018).

10. A. Pickar-Oliver, V. Gough, J. D. Bohning, S. Liu, J. N. Robinson-Hamm, H. Daniels, W. H. Majoros, G. Devlin, A. Asokan, C. A. Gersbach, Full-length Dystrophin Restoration via Targeted Genomic Integration by AAV-CRISPR in a Humanized Mouse Model of Duchenne Muscular Dystrophy. Molecular Therapy (2021), doi:10.1016/j.ymthe.2021.09.003.

11. S. Tong, B. Moyo, C. M. Lee, K. Leong, G. Bao, Engineered materials for in vivo delivery of genome-editing machinery. Nat Rev Mater 4, 726–737 (2019).

12. J. R. Mendell, S. A. Al-Zaidy, L. R. Rodino-Klapac, K. Goodspeed, S. J. Gray, C. N. Kay, S. L. Boye, S. E. Boye, L. A. George, S. Salabarria, M. Corti, B. J. Byrne, J. P. Tremblay, Current Clinical Applications of In Vivo Gene Therapy with AAVs. Molecular Therapy 29, 464–488 (2021).

13. A. Manini, E. Abati, A. Nuredini, S. Corti, G. P. Comi, Adeno-Associated Virus (AAV)-Mediated Gene Therapy for Duchenne Muscular Dystrophy: The Issue of Transgene Persistence. Front Neurol 12, 814174 (2022).

14. C. Li, R. J. Samulski, Engineering adeno-associated virus vectors for gene therapy. Nat Rev Genet 21, 255–272 (2020).

15. K. Y. Chan, M. J. Jang, B. B. Yoo, A. Greenbaum, N. Ravi, W.-L. Wu, L. Sánchez-Guardado, C. Lois, S. K. Mazmanian, B. E. Deverman, V. Gradinaru, Engineered AAVs for efficient noninvasive gene delivery to the central and peripheral nervous systems. Nat. Neurosci. 20, 1172–1179 (2017).

16. U. R. Monani, M. Sendtner, D. D. Coovert, D. W. Parsons, C. Andreassi, T. T. Le, S. Jablonka, B. Schrank, W. Rossoll, W. Rossol, T. W. Prior, G. E. Morris, A. H. Burghes, The human centromeric survival motor neuron gene (SMN2) rescues embryonic lethality in Smn(- /-) mice and results in a mouse with spinal muscular atrophy. Hum Mol Genet 9, 333–339 (2000).

17. T. T. Le, L. T. Pham, M. E. R. Butchbach, H. L. Zhang, U. R. Monani, D. D. Coovert, T. O. Gavrilina, L. Xing, G. J. Bassell, A. H. M. Burghes, SMNDelta7, the major product of the centromeric survival motor neuron (SMN2) gene, extends survival in mice with spinal muscular atrophy and associates with full-length SMN. Hum Mol Genet 14, 845–857 (2005).

18. B. Schrank, R. Götz, J. M. Gunnersen, J. M. Ure, K. V. Toyka, A. G. Smith, M. Sendtner, Inactivation of the survival motor neuron gene, a candidate gene for human spinal muscular atrophy, leads to massive cell death in early mouse embryos. PNAS 94, 9920–9925 (1997).

19. K. Suzuki, M. Yamamoto, R. Hernandez-Benitez, Z. Li, C. Wei, R. D. Soligalla, E. Aizawa, F. Hatanaka, M. Kurita, P. Reddy, A. Ocampo, T. Hishida, M. Sakurai, A. N. Nemeth, E. N. Delicado, J. M. Campistol, P. Magistretti, P. Guillen, C. R. Esteban, J. Gong, Y. Yuan, Y. Gu, G.-H. Liu, C. López-Otín, J. Wu, K. Zhang, J. C. I. Belmonte, Precise in vivo genome editing via single homology arm donor mediated intron-targeting gene integration for genetic disease correction. Cell Res 29, 804–819 (2019).

20. T. Dangouloff, L. Servais, Clinical Evidence Supporting Early Treatment Of Patients With Spinal Muscular Atrophy: Current Perspectives. Ther Clin Risk Manag 15, 1153–1161 (2019).

21. A. Besse, S. Astord, T. Marais, M. Roda, B. Giroux, F.-X. Lejeune, F. Relaix, P. Smeriglio, M. Barkats, M. G. Biferi, AAV9-Mediated Expression of SMN Restricted to Neurons Does Not Rescue the Spinal Muscular Atrophy Phenotype in Mice. Molecular Therapy (2020), doi:10.1016/j.ymthe.2020.05.011.

22. K. Kotulska, A. Fattal-Valevski, J. Haberlova, Recombinant Adeno-Associated Virus Serotype 9 Gene Therapy in Spinal Muscular Atrophy. Front Neurol 12, 726468 (2021).

23. V. Robin, G. Griffith, J.-P. L. Carter, C. J. Leumann, L. Garcia, A. Goyenvalle, Efficient SMN Rescue following Subcutaneous Tricyclo-DNA Antisense Oligonucleotide Treatment. Mol Ther Nucleic Acids 7, 81–89 (2017).

24. H. Zhou, N. Janghra, C. Mitrpant, R. L. Dickinson, K. Anthony, L. Price, I. C. Eperon, S. D. Wilton, J. Morgan, F. Muntoni, A Novel Morpholino Oligomer Targeting ISS-N1 Improves Rescue of Severe Spinal Muscular Atrophy Transgenic Mice. Hum Gene Ther 24, 331–342 (2013).

25. L. Wang, H. Wang, P. Bell, D. McMenamin, J. M. Wilson, Hepatic gene transfer in neonatal mice by adeno-associated virus serotype 8 vector. Hum Gene Ther 23, 533–539 (2012).

26. M.-O. Deguise, Y. D. Repentigny, A. Tierney, A. Beauvais, J. Michaud, L. Chehade, M. Thabet, B. Paul, A. Reilly, S. Gagnon, J.-M. Renaud, R. Kothary, Motor transmission defects with sex differences in a new mouse model of mild spinal muscular atrophy. eBioMedicine 55 (2020), doi:10.1016/j.ebiom.2020.102750.

27. M. D. Howell, E. W. Ottesen, N. N. Singh, R. L. Anderson, R. N. Singh, Gender-Specific Amelioration of SMA Phenotype upon Disruption of a Deep Intronic Structure by an Oligonucleotide. Mol Ther 25, 1328–1341 (2017).

28. M. Arbab, Z. Matuszek, K. M. Kray, A. Du, G. A. Newby, A. J. Blatnik, A. Raguram, M. F. Richter, K. T. Zhao, J. M. Levy, M. W. Shen, W. D. Arnold, D. Wang, J. Xie, G. Gao, A. H. M. Burghes, D. R. Liu, Base editing rescue of spinal muscular atrophy in cells and in mice. Science, eadg6518 (2023).

29. H.-K. Liao, F. Hatanaka, T. Araoka, P. Reddy, M.-Z. Wu, Y. Sui, T. Yamauchi, M. Sakurai, D. D. O’Keefe, E. Núñez-Delicado, P. Guillen, J. M. Campistol, C.-J. Wu, L.-F. Lu, C. R. Esteban, J. C. Izpisua Belmonte, In Vivo Target Gene Activation via CRISPR/Cas9-Mediated Trans-epigenetic Modulation. Cell 171, 1495-1507.e15 (2017).

30. J. I. Ayers, S. Fromholt, O. Sinyavskaya, Z. Siemienski, A. M. Rosario, A. Li, K. W. Crosby, P. E. Cruz, N. M. DiNunno, C. Janus, C. Ceballos-Diaz, D. R. Borchelt, T. E. Golde, P. Chakrabarty, Y. Levites, Widespread and Efficient Transduction of Spinal Cord and Brain Following Neonatal AAV Injection and Potential Disease Modifying Effect in ALS Mice. Molecular Therapy 23, 53–62 (2015).

31. M. Martin, Cutadapt removes adapter sequences from high-throughput sequencing reads. EMBnet.journal 17, 10–12 (2011).

32. N. L. Bray, H. Pimentel, P. Melsted, L. Pachter, Near-optimal probabilistic RNA-seq quantification. Nat Biotechnol 34, 525–527 (2016).

33. Y. Liao, J. Wang, E. J. Jaehnig, Z. Shi, B. Zhang, WebGestalt 2019: gene set analysis toolkit with revamped UIs and APIs. Nucleic Acids Res 47, W199–W205 (2019).

34. Fast and accurate long-read alignment with Burrows–Wheeler transform | Bioinformatics | Oxford Academic (available at https://academic.oup.com/bioinformatics/article/26/5/589/211735).

