## Supplemental Figures and Movies for "Therapeutic strategy for spinal muscular atrophy by combining gene supplementation and genome editing": Sup Figure 4.pdf

### Figure S4

**A**

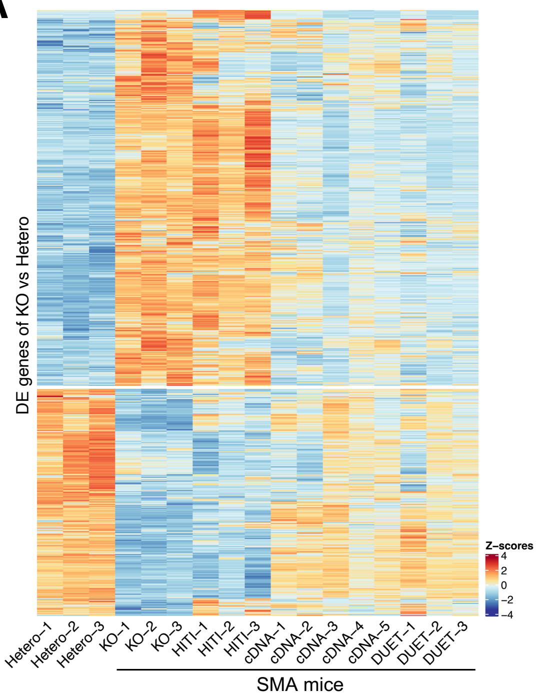


**D**

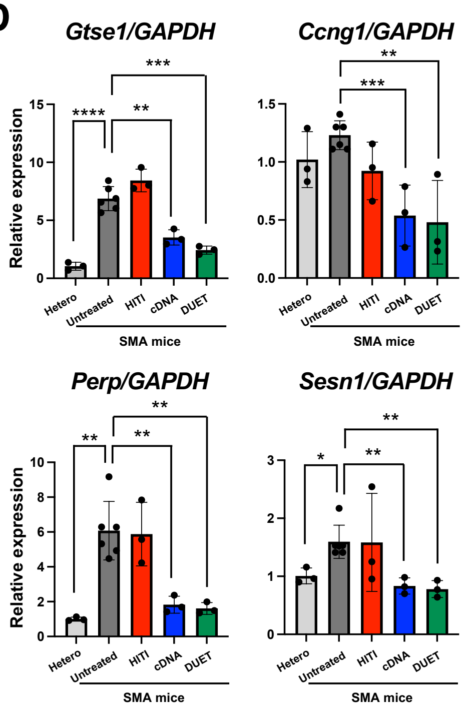


**B** Upregulated biological process in SMA mice (vs Heterozygous mice)

| Description | Genes | *p* value | FDR |
|---|---|---|---|
| drug catabolic process | 38 | 4.37E-14 | 3.55E-11 |
| protein activation cascade | 20 | 2.04E-10 | 8.27E-08 |
| negative regulation of hydrolase activity | 58 | 9.17E-08 | 2.48E-05 |
| organic hydroxy compound metabolic process | 57 | 2.68E-07 | 5.45E-05 |
| regulation of peptidase activity | 56 | 3.43E-07 | 5.57E-05 |
| lipid catabolic process | 41 | 6.93E-07 | 8.03E-05 |
| organic anion transport | 50 | 9.53E-06 | 5.53E-04 |
| response to peptide | 52 | 2.72E-05 | 8.89E-04 |
| response to wounding | 53 | 4.01E-05 | 1.16E-03 |
| circulatory system process | 49 | 1.55E-04 | 3.60E-03 |

**C** Downregulated biological process in SMA mice (vs Heterozygous mice)

| Description | Genes | *p* value | FDR |
|---|---|---|---|
| monovalent inorganic cation transport | 42 | 3.40E-10 | 2.76E-07 |
| regulation of ion transmembrane transport | 36 | 2.36E-08 | 9.59E-06 |
| regulation of membrane potential | 34 | 1.07E-07 | 2.90E-05 |
| muscle system process | 26 | 2.63E-05 | 5.34E-03 |
| receptor-mediated endocytosis | 20 | 5.30E-05 | 8.61E-03 |
| neurotransmitter transport | 21 | 1.23E-04 | 1.43E-02 |
| microtubule-based movement | 19 | 1.68E-04 | 1.52E-02 |
| locomotory behavior | 18 | 1.94E-04 | 1.58E-02 |
| circulatory system process | 28 | 2.43E-04 | 1.80E-02 |
| sensory perception of mechanical stimulus | 14 | 6.74E-04 | 3.65E-02 |
