## Supplementary figures and images for "Therapeutic strategy for spinal muscular atrophy by combining gene supplementation and genome editing"

### Sup Figure 1.pdf

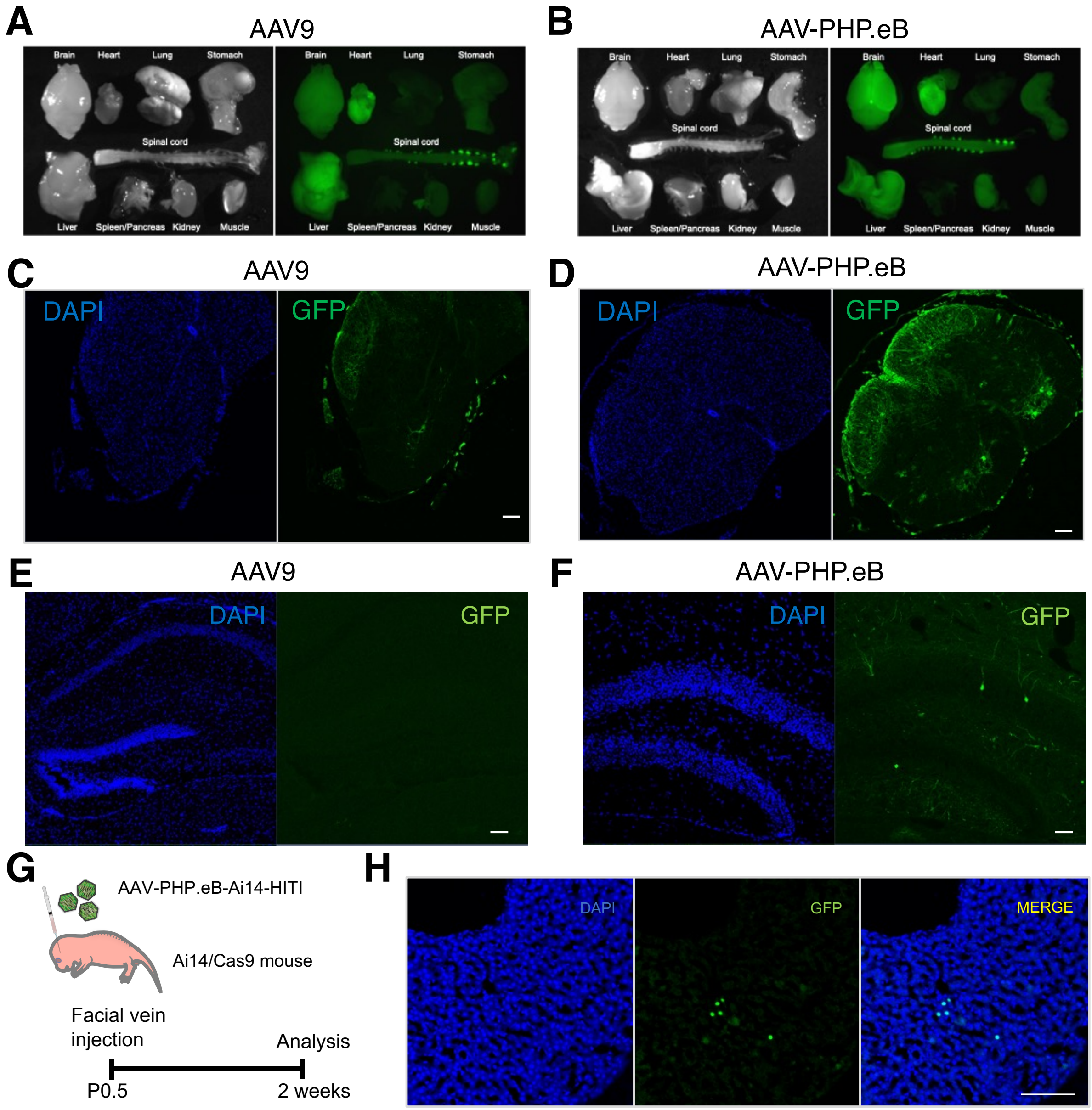

### Sup Figure 2.pdf

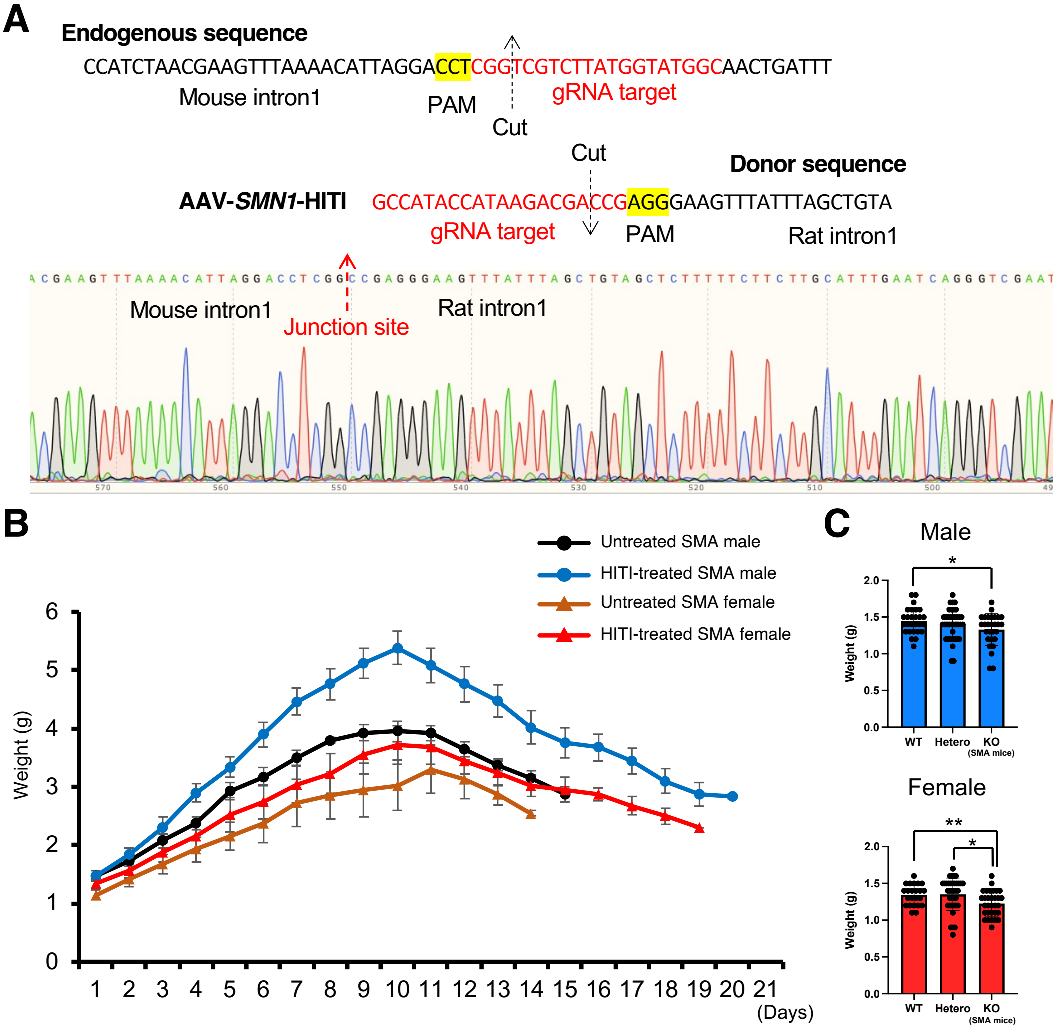

### Sup Figure 3.pdf

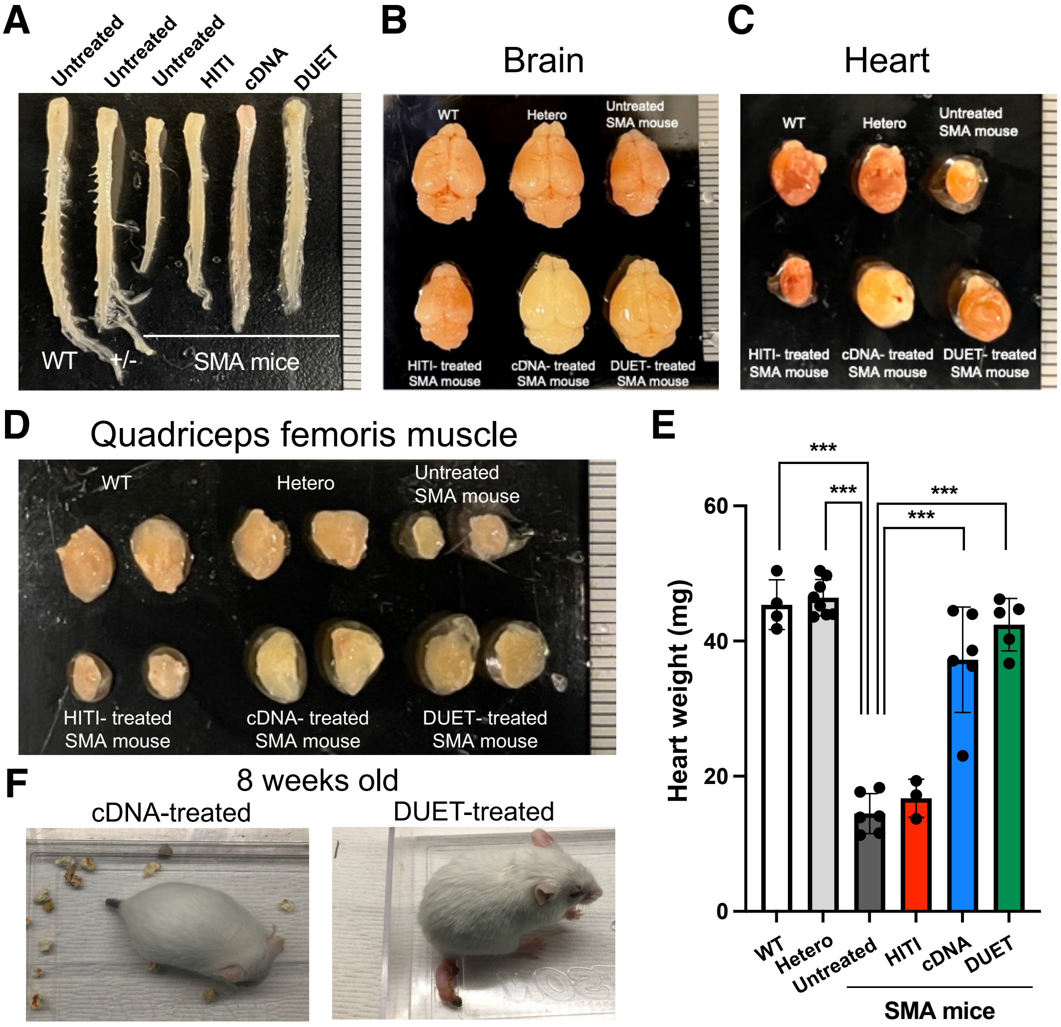

### Sup Figure 5.pdf

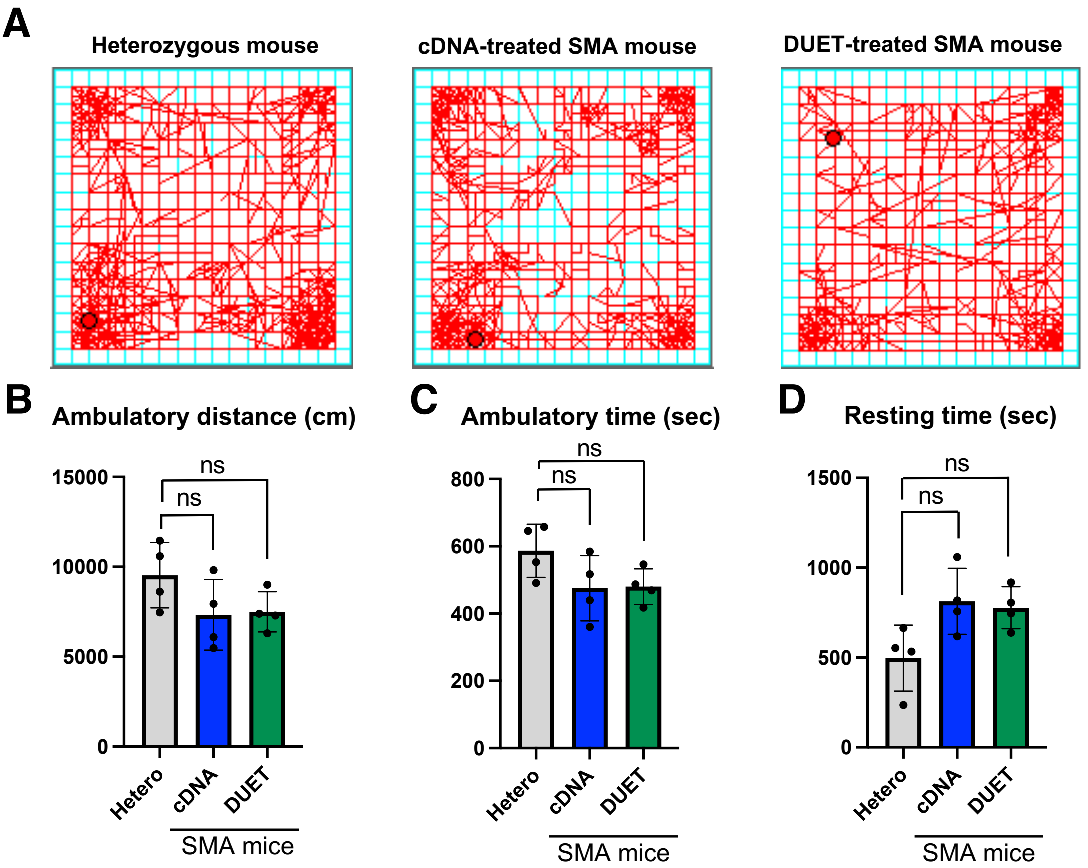

### Sup Figure 6.pdf

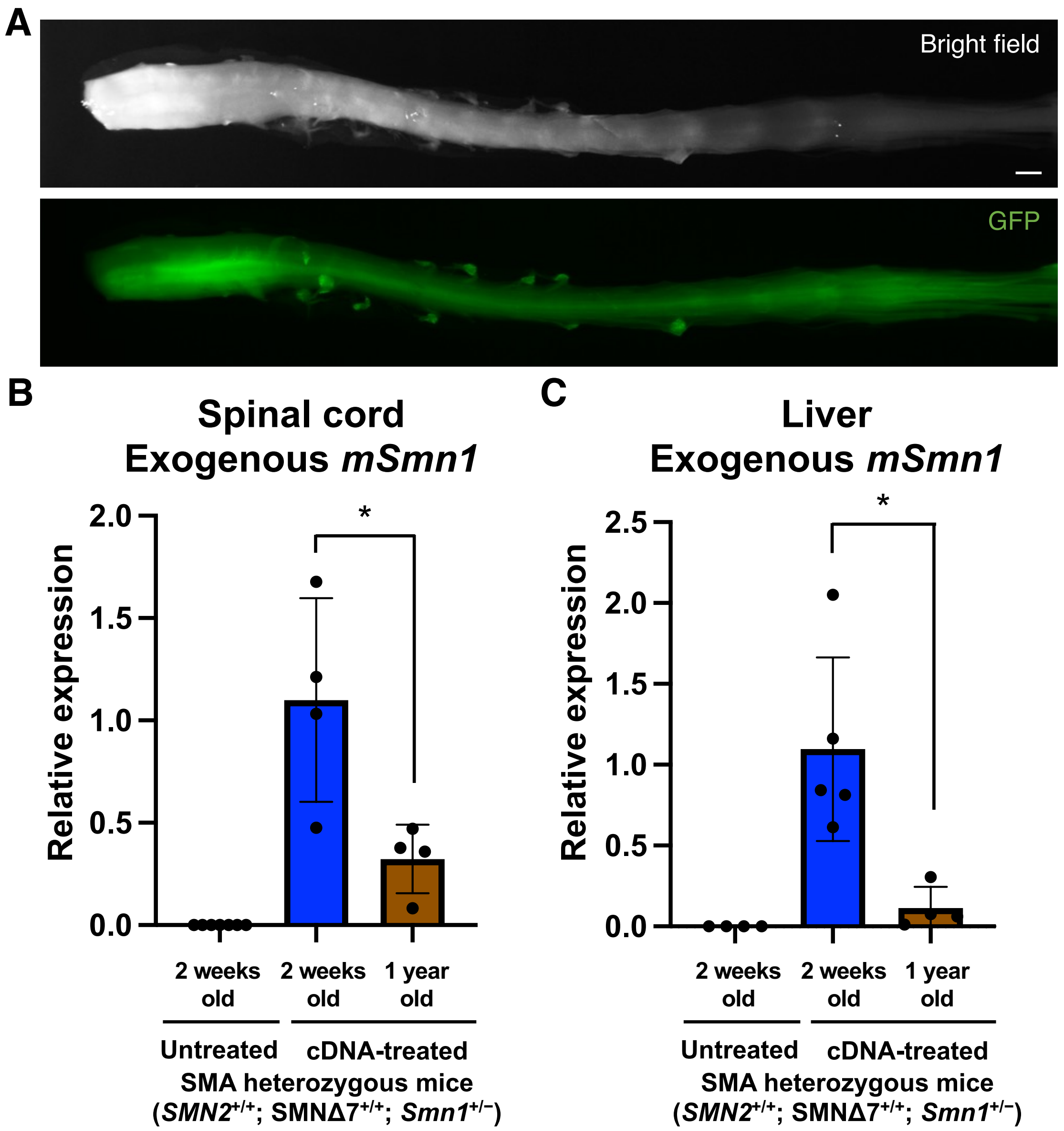
